# Comparative Population Genomics across Three Continents Reveals Immune Adaptation and Implications for Vaccine Development

**DOI:** 10.64898/2026.08.03.741619

**Authors:** Ahmed Tawfik, Kristel Van Steen, Anavaj Sakuntabhai

## Abstract

Human populations have evolved distinct immune responses to local pathogens, and investigating signatures of natural selection across populations can reveal both shared and population-specific immune response mechanisms. We compared genomic signatures of natural selection in Senegalese, Thai, and Peruvian populations using two complementary methods, hypothesising that this tri-continental comparison would reveal population-specific adaptive variants in genes regulating immune pathways. Our findings confirm that genetic diversity among these populations is reflected in their immune responses to regionally endemic pathogens. Furthermore, candidate immune-related genomic regions identified through selection scanning provide insights for ancestry-informed vaccine development by directing antigen design for region-specific pathogens, incorporating population-specific adjuvants, and tailoring administration strategies.

## Introduction

Human populations exhibit significant genomic diversity due to migration, genetic drift, and adaptation to various environmental conditions [1]. Pathogens, including viruses, parasites and enteric bacteria, have exerted selective pressures in the human genome, resulting in detectable signatures of natural selection, particularly within genes involved in immune defence [2–4]. A comparative analysis of these signatures across different populations can reveal both shared and population-specific immune response mechanisms that have developed in response to local pathogens and disease threats [5].

Recently, comparative population genomics has emerged as a systemic approach for detecting signatures of natural selection and local adaptation. An emerging method, such as the Population Branch Statistic (PBS), identifies genomic loci where allele frequencies differ significantly between populations and returns specific variants that may have been affected by selective pressures [6, 7]. In addition, the Integrated Haplotype Score (iHS) method detects recent positive selection within populations by assessing extended haplotype homozygosity surrounding variants undergoing adaptation [8]. Both PBS and iHS collaboratively could establish a robust framework for identifying genetic variants and associated pathways essential to human adaptation to a wide range of ecological conditions, including immune defence against endemic pathogens [9, 10].

Despite these advances in comparative population genomics, several research studies remain focused on single populations or comparisons within a single continent, leaving a gap in targeted analysis of populations from distant geographic regions with unique disease burdens. The populations of Senegal (West Africa), Thailand (Southeast Asia), and Peru (South America) serve as examples of such cohorts, where each has evolved in distinct pathogen environments. For instance, malaria is considered an endemic pathogen that has imposed significant selective pressure across these regions, and populations have developed independent evolutionary responses, as evidenced by population-specific resistance alleles [11]. Consequently, a focused comparison of these three populations facilitates the identification of both shared and population-specific genetic adaptations against malaria and other pathogens.

This study conducts a comparative analysis of Single Nucleotide Polymorphism (SNP) datasets from Senegalese, Thai, and Peruvian populations utilising PBS and iHS methods, hypothesising that this tri-continental comparison will reveal a fundamental set of genes and immune pathways commonly targeted by natural selection, alongside population-specific selections on single-nucleotide variants indicative of localised historical pathogen pressures. This comparative analysis revealed focused adaptations against a wide range of endemic regional pathogens and identified key immune-related genes recurrently targeted by selection across the three populations. These immune-related genes extend beyond peptide processing and antigen presentation, encompassing other facets of the immune response, including adaptations to interferon signalling via distinct molecular mechanisms and consistent selection of immune regulation in conjunction with immune activation, reflecting the importance of the trade-off between immunostimulation and immunomodulation. These evolutionary signatures have direct implications for vaccine development, guiding improved antigen design, population-specific adjuvant choices, and the customisation of delivery platforms [12, 13]. The insights we obtained from integrating comparative population genomics with translational immunology as a part of this study may offer a strategy to advance the development of ancestry-informed vaccines and promote equitable global health.

## Methods

### Subjects and Study Protocol

The project protocol was primarily intended to understand infectious diseases among populations. The objectives were carefully explained to individuals participating from each population, and informed consent was obtained from all subjects or the guardians of children younger than 15 years. The project was initially approved by the Ministry of Health of Senegal and the assembled village population. Approval was then renewed on a yearly basis. Audits were done regularly by the National Ethics Committee of Senegal and ad hoc committees of the Ministry of Health, the Pasteur Institute in Dakar, Senegal, and the Institut de Recherche pour le Développement in Marseille, France.

### Genomic Dataset Preprocessing and Quality Control

Whole-genome SNP genotype data were generated on the Illumina HumanOmniExpress-24v1-0 BeadChip from three population cohorts: Senegalese (West Africa), Thai (Southeast Asia), and Peruvian (South America). Standard quality control (QC) procedures were applied separately for each population using the R environment [14]. Individuals with sex chromosome mismatches and a call rate below 90%, as well as SNPs with a call rate below 90%, were excluded. Autosomal SNPs were imputed, and haplotype phasing was estimated using Beagle 5.5 [15, 16]. Following quality control, datasets from the three cohorts were merged, and only autosomal SNPs in all three populations were retained to ensure comparable genomic profiles for subsequent analysis. The final quality-controlled dataset comprised 685,589 autosomal SNPs per individual and 1,212 individuals: 404 from Senegal, 404 from Thailand, and 404 from Peru.

### Population Genetic Diversity Assessment

To confirm the genetic distinctiveness of the three populations and validate their suitability for comparative genome-wide natural selection scans, two complementary analyses were conducted using genome-wide SNP data.

- **Principal Component Analysis (PCA)** is an unsupervised dimensionality reduction technique that transforms a dataset of correlated SNPs into a minimal set of uncorrelated orthogonal principal components (PCs). The first two PCs capture the largest sources of genetic variation, enabling visualisation of genome-wide SNP data in two dimensions and revealing the structure of the three populations. PCA was performed using the snpgdsPCA function in the SNPRelate R package[17].
- **Hierarchical Clustering** constructs a dendrogram of samples based on pairwise genetic distances in an unsupervised manner without assuming predefined groups. A pairwise genetic distance matrix was computed using the identity-by-state (IBS) method via the snpgdsIBS function in the SNPRelate R package. The snpgdsIBS function calculates the fraction of identity-by-state for each pair of individuals, providing a robust genetic distance metric for population structure analysis. The resulting matrix was subjected to hierarchical clustering in SNPRelate to visualise the dendrogram and confirm the separation of individuals into three major clusters corresponding to the Senegalese, Thai, and Peruvian populations.

### Linkage Disequilibrium Calculation and Decay Estimation

Linkage disequilibrium (LD) is defined as the non-random association of alleles at different loci [18], where certain allele correlations occur, reflecting patterns of recombination rates. LD is quantified using the squared correlation coefficient (r²) between pairs of SNPs [19], and the coefficients of r² typically decline with the increase of genetic distance between loci along a chromosome, resulting in LD decay [20,21]. To characterise LD decay across the genome, the mean r² between SNP pairs was assessed as a function of physical distance in kilobases. For each population, all SNP pairs were binned by distance into six intervals (0–10 kb, 10–50 kb, 50–100 kb, 100–200 kb, 200–500 kb, and 500–1000 kb). The mean r² was calculated per bin, representing the average non-random association of alleles within each distance bin. The distance at which mean r² dropped below 0.1 was defined as the LD decay threshold. LD estimation and r² calculation were performed separately for each population using PLINK v1.9 [22].

### Natural Selection Signatures Detection

Natural selection in humans is the process by which environmental pressures favour individuals with advantageous genetic traits, thereby improving their survival and reproductive success [23]. To capture signals of natural selection in the three populations, two complementary methods were employed:

**1-Population Branch Statistic (PBS)** quantifies population-specific allele frequency divergence in comparison to two reference populations along a trifurcating tree. PBS identifies SNPs whose allele frequencies have changed in that population alone, indicating that the population has adapted over extended periods of time. To compute PBS, pairwise fixation indices (Fst) for each SNP between all three population pairs (Fst(Senegal, Thailand), Fst(Senegal, Peru), and Fst(Thailand, Peru)) were calculated using the snpgdsFst function in the SNPRelate R package according to the Weir & Cockerham estimation method [24]. The branch length metric for the three population pairs is a log-transformed function of Fst, which accumulates additively with time and can be calculated for each population pair as follows:

T(Senegal, Thailand) = -log(1-Fst(Senegal, Thailand))

T(Senegal, Peru) = -log(1-Fst(Senegal, Peru))

T(Thailand, Peru) = -log(1-Fst(Thailand, Peru))

Finally, PBS was estimated for each population by isolating the population-specific branch length as follows:

PBS(Senegal) = (T(Senegal, Thailand) + T(Senegal, Peru) - T(Thailand, Peru)) / 2

PBS(Thailand) = (T(Senegal, Thailand) + T(Thailand, Peru) - T(Senegal, Peru)) / 2

PBS(Peru) = (T(Senegal, Peru) + T(Thailand, Peru) - T(Senegal, Thailand)) / 2

Under neutral conditions, PBS values are typically less than 0.3 in any population, and values exceeding 0.5 are considered suitable candidates for a selection scan. Consistent with established practice in the field, SNPs with PBS > 0.5 were selected as candidates for functional annotation, representing loci in the upper tail of the genome-wide divergence distribution where population-specific allele frequency changes are most noticeable and are considered potential candidates.

**2-Integrated Haplotype Score (iHS)** detects recent or ongoing selection by comparing extended haplotype homozygosity around a core SNP between ancestral and derived alleles. iHS measures relative haplotype homozygosity around a DNA variant, comparing stretches of identical DNA for the ancestral versus derived alleles. Identification of long haplotypes carrying the derived allele at high frequency indicates a recent selective sweep and results in higher absolute iHS values. The calculated score captures ongoing or recent adaptive events that have not yet reached fixation by examining an extended haplotype structure, making it highly complementary to PBS, which captures long evolutionary adaptive events. iHS was estimated within each population using the rehh R package on phased genotype data [25].

Under neutral conditions, iHS values in any population follow a standard normal distribution centred around 0, with most values ranging between -2 and +2 due to typical haplotype decay patterns from neutral recombination events, and values exceeding |iHS| > 2 possess relatively long haplotypes around one allele within that population, consistent with recent positive selection. SNPs with |iHS| > 2 were selected as candidates for functional annotation, representing the standard empirical threshold applied in genome-wide selection scans to identify loci where extended haplotype homozygosity is inconsistent with neutral expectations.

### Functional Annotation and Impact Prediction

To identify independent genomic loci under selection and avoid redundancy from linked SNPs, an LD-based approach was applied to each population. First, genome-wide pairwise r² values were calculated, and an LD block was defined as a contiguous set of SNPs where all genome-wide pairwise r² values between SNPs within the block were ≥ 0.8. Blocks were constructed using a greedy clustering algorithm, starting with the SNP with the highest PBS or |iHS| value as a tag SNP. Then, all SNPs with r² ≥ 0.8 were merged into a block. Next, blocks that did not include any candidate SNPs with PBS > 0.5 or |iHS| > 2 after merging were discarded. This ensured that each LD block reported either a signal of divergent selection with PBS > 0.5 or a signal of recent selection with |iHS| > 2 at an independent genomic locus. Finally, all PBS and iHS tag SNPs, and all other SNPs in LD within each LD block were annotated using the Ensembl Variant Effect Predictor (VEP) [26]. VEP identifies overlapping transcripts with SNPs and uses a rule-based approach to predict the effects that each variant allele may have on each transcript by mapping different types of genetic variants onto a gene structure from the 5’ end to the 3’ end, which include:

- **Upstream gene variants** are sequence variants located near the 5’ end of a gene that may affect promoter regions or other regulatory elements controlling transcription [27].
- **5 prime UTR variants** are sequence variants located in the 5′ untranslated region (5′ UTR) of a gene, and changes in this region can influence protein translation [28].
- **Stop-gained variants** introduce premature termination codons, leading to shortened or nonfunctional proteins and often resulting in loss of gene function [29].
- **Missense variants** are amino acid substitutions in a protein’s sequence that can affect its function [30].
- **Synonymous variants** are nucleotide changes that result in no amino acid substitutions but may affect splicing or regulatory elements [31].
- **Splice donor region variants** are sequence variants that fall in the gene’s 5’ splice site at exon-intron boundaries and may affect the splicing process [32].
- **Splice polypyrimidine tract variants** are sequence variants that fall in the polypyrimidine tract, a part of the splice acceptor site that may impact mRNA splicing [32].
- **Intron variants** are sequence variants occurring within an intron that result in no amino acid substitutions but may affect splicing or regulatory elements [33].
- **3 prime UTR variants** are sequence variants located in the 3′ untranslated region (3′ UTR) of a gene, and changes in this region may affect mRNA stability or miRNA binding [28].
- **Downstream gene variants** are sequence variants located near the 3’ end of a gene that may affect the distal regulatory regions [34].

We utilised VEP to map the candidate SNPs to their target genes and retrieve their corresponding Gene Ontology (GO) terms [35]. All variants within a naturally selected LD block were considered potentially relevant to the adaptive signal, and their effects on immune-related genes were interpreted using GO terms associated with the gene transcripts. Since natural selection acts on haplotypes, any SNP in strong linkage disequilibrium (r² ≥ 0.8) with a significant tag SNP may represent the true causal variant or contribute to the selected phenotype through joint effects. Variants within each block, including intronic, UTR, and intergenic SNPs, were included because they may exert regulatory effects, alter splicing, or be in LD with untyped causal mutations (Figure 1). This comprehensive approach captures both coding and non-coding adaptive variations, reflecting the multifaceted nature of selection on immune responses.

**Figure 1:**
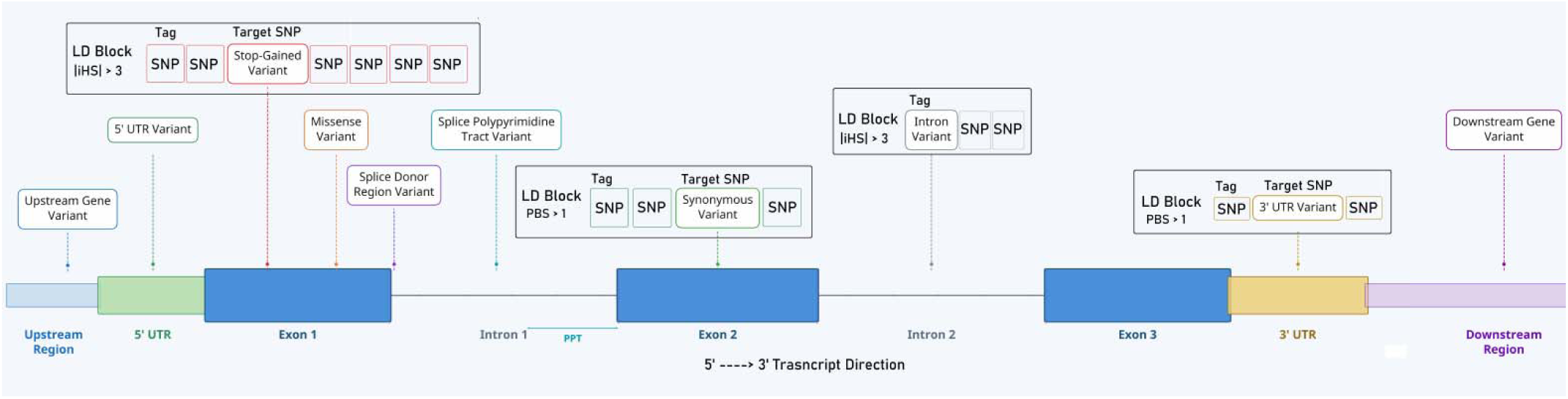
Schematic example for a gene’s adaptation scenario. Exon 1 includes a stop-gained variant within an LD block tagged by a SNP with |iHS| > 2, reflecting recent selection. Exon 2 includes a synonymous variant within an LD block tagged by a SNP with PBS > 0.5, reflecting divergent selection. Intron 2 includes an intronic Tag SNP with |iHS| > 2. The 3′ untranslated region includes a 3’ UTR variant as part of an LD block tagged with a SNP with PBS > 0.5.

## Results

### Population-Specific LD Patterns in Senegalese, Thai, and Peruvian Populations

The Senegalese, Thai, and Peruvian populations are geographically and historically distinct human groups. Their genetic divergence results from different ancient migration routes, varying effective population sizes, and exposure to unique selective pressures, including local pathogens [36]. To better reveal and understand these differences, we assessed genetic differentiation among these populations using PCA and hierarchical clustering of genome-wide SNP data; both methods demonstrated clear separation by geographic origin and confirmed substantial genetic divergence (Figure 2). Notably, hierarchical clustering reveals within-population diversity based on pairwise genetic dissimilarity between individuals, which suggests that Senegal has the highest within-population diversity, while Peru has the lowest. This diverse genetic structure between populations and within each population indicates that the three populations have experienced independent evolutionary histories, affected by different demographic events and local selective pressures. The genetic distinctiveness of these populations suggests studying LD patterns within each population to obtain preliminary insights into how ecological conditions have differently affected their genomes [37].

**Figure 2:**
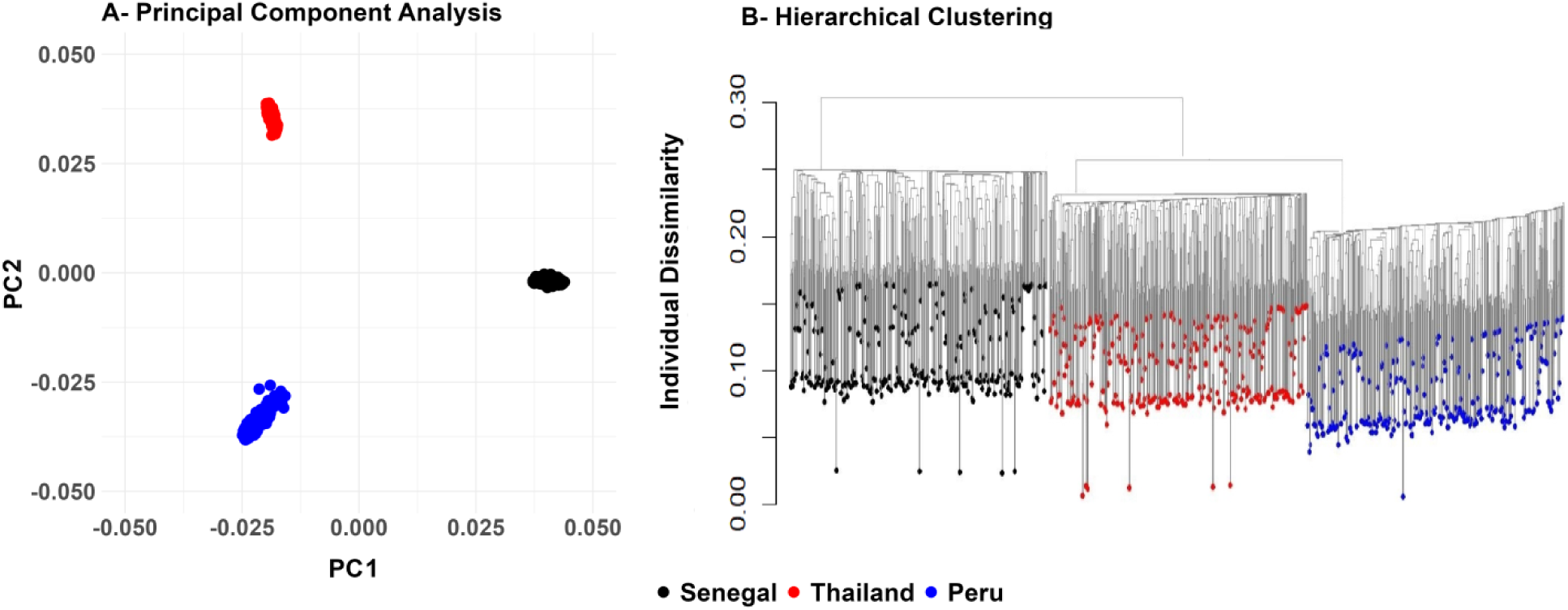
A- Principal component analysis shows three distinct clusters for the three populations along the first two principal components. B- Hierarchical clustering forms three subclusters. The Senegalese cluster is separate from the Thai and Peruvian pair, indicating that the Senegal samples are the most genetically distinct. The vertical height reflects within-population dissimilarity, with Senegal samples showing the greatest vertical variation among the three populations.

LD in finite-sized populations refers to the non-random association of alleles at different loci. The magnitude of these allele associations generally decreases with increasing physical genomic distance between loci, resulting in LD decay. The rate of LD decay is inversely proportional to the effective population size, as larger populations experience higher recombination rates, resulting in a faster approach to linkage equilibrium. To study the LD patterns in these three populations, we calculated pairwise r² values for SNPs within 1 Mb windows, by grouping genomic distances into six bins (0–10 kb, 10–50 kb, 50–100 kb, 100–200 kb, 200–500 kb, and 500–1000 kb), and for each population, we calculated the number of SNP pairs in LD per bin and the mean of r² per bin (Table 1). The Senegalese population had the lowest LD and the fastest LD decay, with an average r² of 0.18 in the 0–10 kb range and a rapid drop to 0.1 at 26 kb. This is consistent with a large effective population size and the absence of major bottlenecks. The Thai population had a higher average r² of 0.32 in the 0–10 kb range and a slower drop to 0.1 at 71 kb. The Peruvian population had the highest average r² of 0.37 in the 0–10 kb range and the slowest drop to 0.1 at 81 kb (Figure 3A). An exception to the overall LD decay trend was observed when comparing the Thai and Peruvian populations (Figure 3B); Thailand exhibited consistently lower mean r² than Peru at shorter genomic distances (<100 kb), but this relationship was reversed in the three longest bins (>100 kb). The higher LD at short distances in Peru is consistent with a reduction in effective population size or a demographic event of admixture, which may increase short-range LD driven by allele-frequency differences between ancestral populations [38]. The LD pattern in Thailand is consistent with East Asian demographic history, which includes the out-of-Africa bottleneck followed by subsequent population expansion, expected to result in a moderate effective population size and slower LD decay at longer genomic distances [39]. Notably, the Senegalese population showed a higher number of outlier LD SNP pairs with elevated r² values in the shortest 0–10 kb bin than the Thai and Peruvian populations (Figure 3A). This LD pattern observed among the Senegalese population and characterised by generally low LD coexisting with an excess of high-LD outlier pairs, could be explained by localised forces that increase LD at specific loci while the remaining genome undergoes rapid recombination. A localised natural selection at certain immune-related genes under local pathogen pressure could account for this pattern, and such regions could be further evaluated for natural selection using PBS and iHS to identify functional variants and haplotype signatures of pathogen-driven adaptation affecting immune-related genes [40].

**Figure 3:**
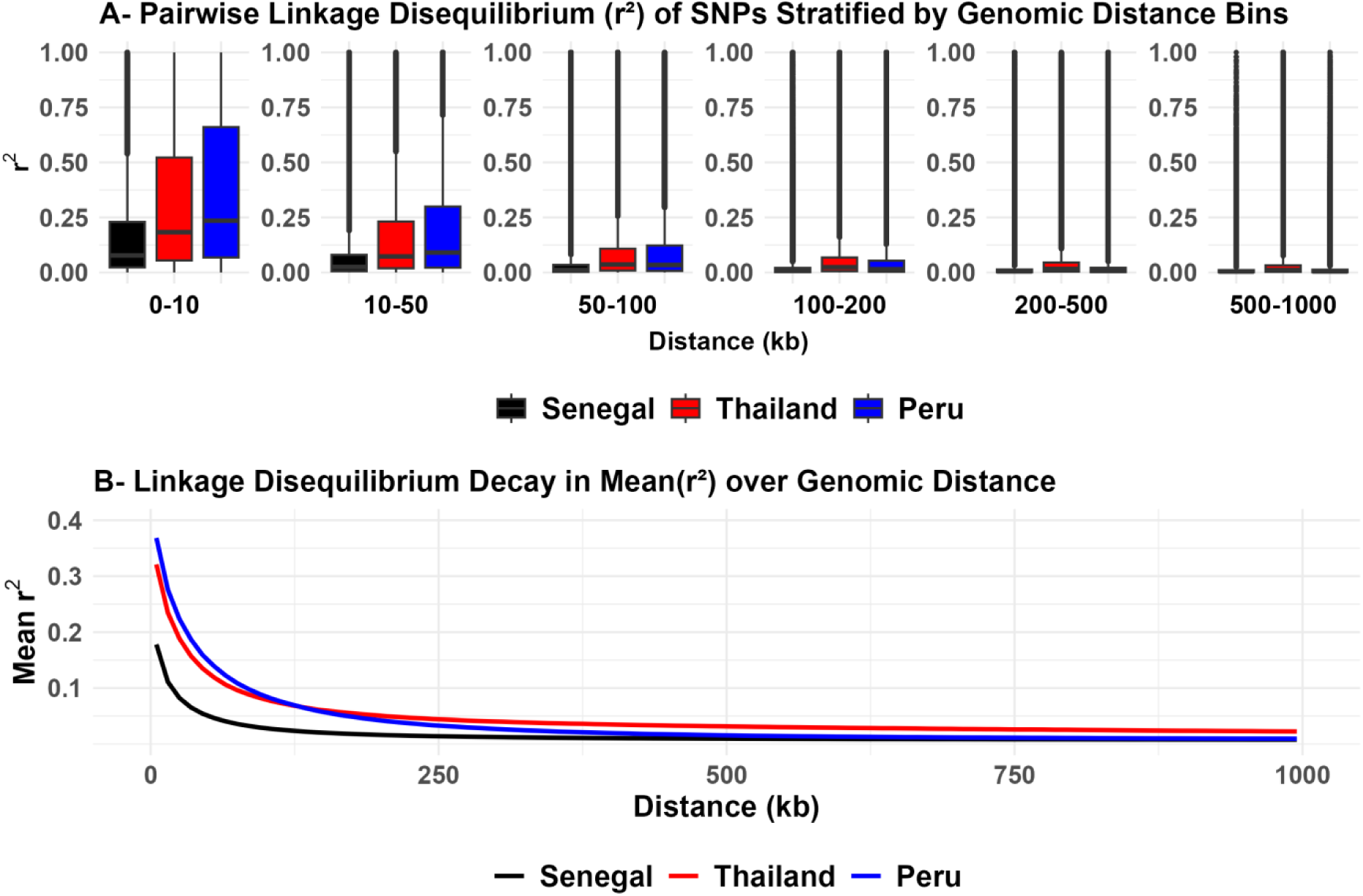
A. Pairwise r² values for SNPs within six distance bins for three populations. B. LD decay curves show mean r² for each bin.

**Table 1:** Number of SNP pairs in LD and mean r² values for SNPs within six distance bins for three populations.

| Bin | Senegal |  | Thailand |  | Peru |  |
| --- | --- | --- | --- | --- | --- | --- |
| | SNP Pairs | mean $r^2$ | SNP Pairs | mean $r^2$ | SNP Pairs | mean $r^2$ |
| **0-10 (kb)** | 1749005 | 0.18 | 1414812 | 0.32 | 1225333 | 0.37 |
| **10-50 (kb)** | 6422151 | 0.08 | 5119195 | 0.18 | 4415017 | 0.21 |
| **50-100 (kb)** | 7636262 | 0.03 | 6028961 | 0.09 | 5184504 | 0.11 |
| **100-200 (kb)** | 14706590 | 0.02 | 11548923 | 0.06 | 9912601 | 0.06 |
| **200-500 (kb)** | 42330359 | 0.01 | 32980806 | 0.03 | 28266700 | 0.02 |
| **500-1000 (kb)** | 68604371 | 0.00 | 53244779 | 0.02 | 45598619 | 0.01 |

### PBS-Derived Signature of Pathogen-Driven Immune Adaptation in Senegal

A comprehensive analysis of the PBS-derived genetic signature in Senegal reveals a cohesive evolutionary strategy influenced by sustained historical pathogen pressure. This strategy consists of three primary components: specialised malaria defence, enhanced antibacterial preparedness, and an integrated system for immunoregulatory balance.

Malaria is one of the oldest diseases in the region [41], and defence mechanisms against it are believed to have co-evolved with human ancestors in Africa. Genomic analyses estimate that the most recent common ancestor of Plasmodium falciparum appeared 50,000 to 100,000 years ago, coinciding with the expansion of Homo sapiens across Africa [42]. In West Africa, Plasmodium falciparum malaria poses a significant health challenge due to its disruption of erythrocyte production through multiple mechanisms, which contributes to severe malarial anaemia (SMA) [43]. As a result, the process of erythrocyte development has been shaped by natural selection in malaria-endemic regions, transforming hematopoietic stem cells (HSCs) into mature red blood cells (RBCs) through sequential lineage commitment, expansion, and maturation. This process balances rapid compensation for anaemia with the risk of parasites exploiting erythroid traits. The Senegalese genome reflects this adaptation, as functional annotation of naturally selected genetic variants includes the molecular process of erythrocyte development (GO:0048821), associated with an intronic variant (rs7833494) in the **ANK1** gene. Variants in ANK1 affect erythrocyte membrane stability and resilience, potentially conferring resistance to Plasmodium falciparum malaria by impeding parasite invasion and maturation or by facilitating the clearance of infected red blood cells [44]. Local adaptation of erythrocyte differentiation is further indicated by annotation of the molecular processes of erythrocyte differentiation (GO:0030218) and regulation of definitive erythrocyte differentiation (GO:0010724). These processes involve the upstream variant (rs1573708) in the **RBFOX2** gene and the intronic variant (rs12262584) in the **EXOC6** gene. The RBFOX2 gene is essential for erythroid differentiation, primarily by regulating the alternative splicing of key cytoskeletal genes required for the mature red blood cell membrane [45,46]. Additionally, a strongly selected downstream variant (rs4669330) in the **ID2** gene and an intronic variant (rs1935949) in the **FOXO3** gene have been under natural selection. ID2 and FOXO3 act as master regulators within the erythrocyte differentiation pathway. FOXO3 is a central transcription factor that promotes erythroid progenitor survival under oxidative stress, a condition intensified by malaria-induced hemolysis, and enhances the longevity of immunological memory cells. This pattern suggests an evolutionary strategy to maintain both erythropoiesis and protective immunity during recurrent infection [47]. ID2 serves as a lineage-fate determinant, favouring erythroid over myeloid differentiation, which may enhance bone marrow output to counter malaria-driven anaemia and support recovery [48]. Concurrently, the humoral component of antiparasitic defence is highlighted by strong signals in complement activation through the classical and lectin pathways (GO:0006958, GO:0001867), regulated by a 3’UTR variant (rs3852053) in the **MASP1** gene, and two variants in the C4b-binding protein genes, a splice donor region variant (rs6690037) in **C4BPB** and an intronic variant (rs7524207) in **C4BPA**. The complement system is a well-established effector mechanism against malaria, mediating the opsonisation and lysis of infected red blood cells, and genetic variation in these regulators modulates disease severity [49]. Complement Receptor 1 variants protect against severe Plasmodium falciparum rosetting, which could explain the selection of the synonymous variant (rs17047661) in the **CR1** gene [50]. The connection between innate recognition and clearance is further emphasised by the annotation of the Fc-gamma receptor signalling pathway involved in phagocytosis (GO:0038096), involving the strongly selected synonymous variant (rs749338) in the **ITPR3** gene. In addition to the molecular processes termed positive regulation of macrophage activation (GO:0043032) and positive regulation of macrophage cytokine production (GO:0060907), involving the missense variant (rs4988956) in the **IL1RL1** modulator gene and the intronic variant (rs304797) in the **CD36** gene, while it is known that CD36 in macrophages enables nonopsonic phagocytosis of infected red blood cells [51]. On the other hand, we identified a synonymous variant (rs12075) in the **DARC** gene, also known as Duffy antigen/receptor for chemokines, which encodes the erythrocyte surface receptor exploited by Plasmodium vivax for host cell invasion. DARC is among the most well-characterised genes associated with malaria resistance mutations in human populations, with multiple functionally relevant variants documented across its coding and regulatory regions [52,53]. The selection of this variant in the Senegalese population is consistent with ongoing evolutionary pressure on this invasion receptor, reflecting the broader pattern of DARC as a recurrent target of pathogen-driven adaptation in malaria-endemic regions.

A similar robust adaptation is evident against invasive bacterial diseases endemic to the region, particularly tuberculosis and leprosy [54]. Adaptation to these bacterial diseases is demonstrated by pervasive selection across the toll-like receptor (TLR) signalling pathway (GO:0002224) and its specific branches, involving the upstream variant (rs10759930) in **TLR4** and the 3’UTR variant (rs3740961) in the downstream adaptor **TRAF6**. The TRAF6 gene is a key adaptor in the TLR signalling pathway, functioning as an E3 ubiquitin ligase downstream of TLRs, including TLR4, to activate NF-κB and MAPK pathways during bacterial recognition. Functional studies indicate that TRAF6 upregulation and interactions enhance bacterial clearance in tuberculosis models [55]. The need for rapid containment of bacteremia is further supported by GO terms for neutrophil chemotaxis (GO:0030593) and leukocyte migration (GO:0050900), which involve the missense variant (rs6135) in the **SELP** gene and the intronic variant (rs9937837) in the **ITGAM** gene. SELP encodes P-selectin, a key adhesion receptor that mediates early steps of leukocyte recruitment and thereby contributes directly to leukocyte migration [56], while ITGAM is a critical integrin for phagocyte adhesion and migration to infection sites [57]. In addition to selecting the missense variant (rs878756) in the **MKL1** gene, which supports host defence against bacteria by maintaining cytoskeletal integrity for pathogen engulfment and immune signalling, its disruption underlies neutrophil defects and increased susceptibility to bacterial infections [58,59].

The evolutionary necessity to elicit strong, recurrent inflammatory responses against persistent pathogens inherently increases the risk of immunopathology, septic shock, and autoimmune damage. This challenge likely drove the emergence of a third pillar of adaptation to maintain immunoregulatory balance and attenuate excessive immune activation. Immune regulation is evident at the level of the innate immune response, with selection of a missense variant (rs2074158) in **DHX58**, also known as LGP2, a negative regulator of RIG-I-like receptor signalling (GO:0039536) that limits excessive interferon production [60]. Additionally, control of lymphocyte dynamics through negative regulation of T cell activation (GO:0050868) is mediated by a 5’UTR variant (rs1039524) in the **TIGIT** gene and a 3’UTR variant (rs3811021) in the **PTPN22** gene. TIGIT is a major co-inhibitory receptor expressed on T cells that suppresses excessive immune responses and has been shown to limit tissue pathology in viral infections [61], while population-specific variants of PTPN22 suggest adaptation to balance infection defence and limit autoimmunity risk [62]. Furthermore, a 3’UTR variant (rs1217382) in the **BCL2L15** gene, predicted to be involved in the apoptotic process, and a missense variant (rs1056932) in the master regulator **BCL6** are implicated in the negative regulation of the type 2 immune response (GO:0002829) and in the regulation of germinal centre formation (GO:0002634). BCL6 is pivotal for T follicular helper cell differentiation and for constraining inflammatory cytokine production, thereby modulating the magnitude and quality of antibody responses [63,64]. Two high-impact stop-gained variants were also identified; one in the neuronal calcium-binding gene **PREP** (rs9486069) and another in the **NECAB2** gene (rs925331), suggesting adaptation through altered neuroendocrine and inflammatory signalling. Loss of NECAB2 function may exert an anti-inflammatory effect in neurogenic pain contexts, functioning as a form of neuroimmune modulation that dampens sensory hypersensitivity and local cytokine responses [65], and provides a selective advantage against infections that preferentially invade peripheral nerves such as leprosy [66]. Conversely, a truncated PREP may lead to the accumulation of immunomodulatory neuropeptides, such as substance P [67], potentially enhancing prompt innate immune defence against endemic infections [68]. Elevated levels of substance P would result in a faster, stronger, and more aggressive innate immune response against nerve pathogens, such as leprosy. Finally, a strongly selected variant (rs916334) was identified in the 5’ untranslated region (5’UTR) of the **APOL3** gene. APOL3 is a documented example of geographical adaptation in West African populations due to its role in innate immunity against trypanosomes, parasitic infections that affect humans and animals and cause diseases such as African sleeping sickness [69].

**Figure 4:**
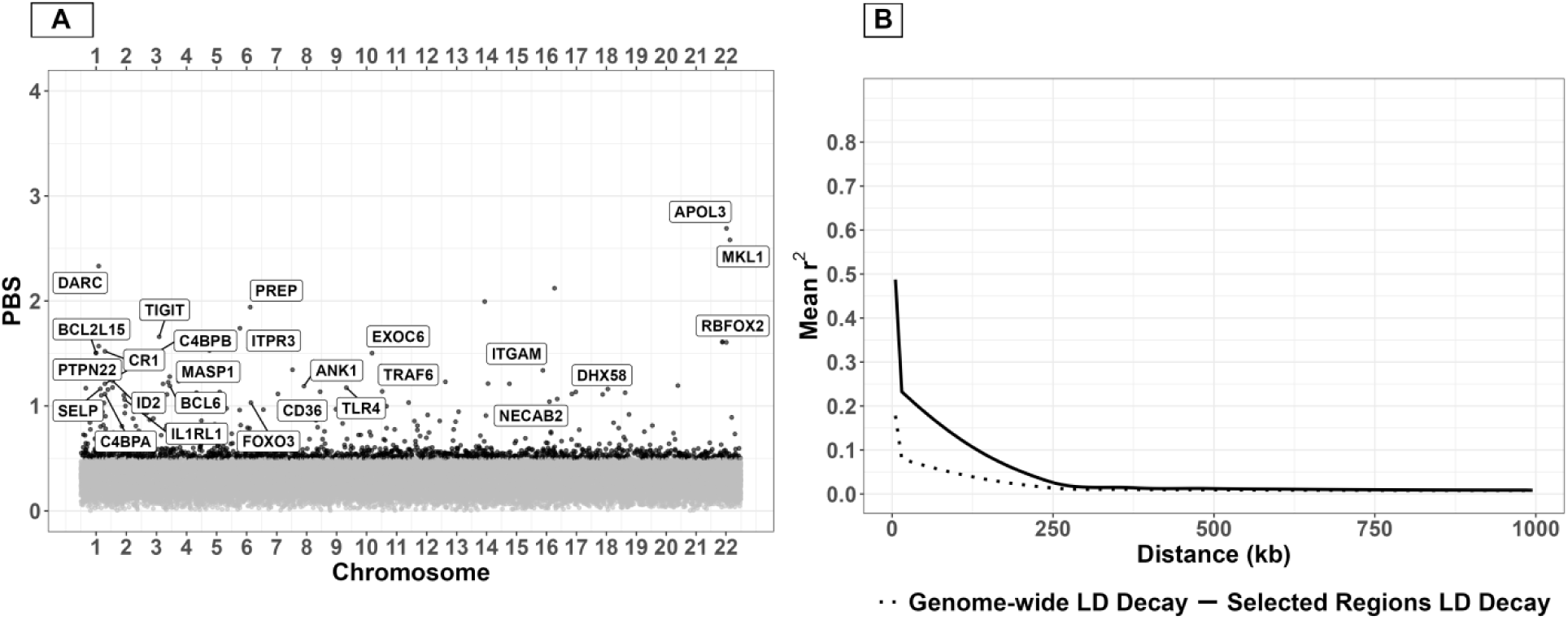
A- Candidate genes under natural selection, identified by genetic variants with elevated PBS values in the Senegalese population. B-LD decay curve for PBS-selected regions is shown in comparison to the genome-wide LD decay curve. The selected regions’ solid curve shows a remarkable difference in mean r² at short distances and a slower decay than the dotted genome-wide decay. The elevated LD in the selected regions reflects genuine selective sweeps.

### iHS-Derived Signature of Pathogen-Driven Immune Adaptation in Senegal

Analysis of recent positive selection using iHS in the Senegalese population reveals a shift in hematopoietic regulation compared to PBS analysis. While PBS indicates ancient selection favouring erythropoiesis via FOXO3 and ID2 for malaria adaptation, iHS demonstrates strong recent selection on an upstream variant (rs7943006) in **LMO2,** a negative regulator of erythrocyte differentiation at later stages. LMO2 is a transcription regulator essential for early erythropoiesis, forming multimeric transcription factor complexes to drive erythroid progenitor proliferation and survival [70]. Naturally selected LMO2 variants may enhance this proliferative capacity, enabling rapid replenishment of red blood cells lost to parasite-induced hemolysis and dyserythropoiesis in malaria anaemia. This malaria resistance gene may represent an evolutionary trade-off or reprioritisation. Another evolutionary trade-off associated with bacterial resistance is indicated by the intronic variant (rs3134930) in **NOTCH4**, since variants that lower NOTCH4 expression may have been favoured for enhancing anti-mycobacterial tuberculosis inflammatory responses [71].

The majority of the iHS signal indicates selection favouring an antiviral immune response and the clearance of endemic viral infections, such as Hepatitis B Virus (HBV), with a carrier rate exceeding 10% in Senegal. Endemic HBV exerts chronic pressure on MHC-I-restricted immune surveillance [72]. Similarly, Lassa virus, endemic to West Africa, requires effective cytotoxic T lymphocyte (CTL) responses for clearance [73], and prevalent herpesviruses necessitate lifelong CD8+ T-cell surveillance, providing a persistent selective background [74]. Rift Valley fever virus (RVF), a zoonotic phlebovirus hyperendemic to the Sahel region, including Senegal’s river basins, causes recurrent outbreaks with significant morbidity and mortality. Clearance of RVF and survival from severe disease depend strongly on CD4+ T cells and virus-specific antibodies [75]. The collective burden of these viruses has produced a genomic signature centred on optimising antigen presentation, CTL activity, and virus-specific antibody responses. Notably, a synonymous variant (rs1800454) in the **TAP2** gene is under selection, which participates in multiple immune molecular processes such as antigen processing and presentation via MHC class I (GO:0002474) and positive regulation of T cell-mediated cytotoxicity (GO:0001916), along with a downstream variant (rs9391846) in **HLA-B** and an upstream variant (rs5010528) in **HLA-C**. Concurrently, the intracellular peptide-loading machinery demonstrates significant recent selection, highlighted by intronic variants in **ERAP1** (rs998509) and **TAPBP** (rs9469397). ERAP1 trims antigenic peptides in the endoplasmic reticulum for optimal MHC-I loading [76], while TAPBP (tapasin) acts as a scaffold in the peptide-loading complex to ensure stable peptide–MHC-I assembly and presentation [77, 78]. Furthermore, the missense variant (rs17587) in **PSMB9** supports the hypothesis that immunoproteasomes containing PSMB9 improve CD8+ T cell recognition of infected or transformed cells [79, 80]. Downstream of antigen presentation, the selection signature extends to effector cell recruitment. This is evidenced by the selection of the chemokine **XCL1**, which involves the 3’UTR variant (rs982143), highlighting its role as a critical regulator of dendritic cell recruitment and CTL priming [81]. This results in the annotation of processes such as positive regulation of T cell chemotaxis (GO:0010820) and mature natural killer cell chemotaxis (GO:0035782). Together, these patterns indicate a comprehensive evolutionary strategy to accelerate the detection and clearance of intracellular viral pathogens. The strategy is further supported by co-selection of the interferon gamma-mediated signalling pathway (GO:0060333), involving the missense variant (rs9666607) in **CD44**. Interferon gamma (IFN-γ), via its JAK-STAT signalling pathway, transcriptionally upregulates HLA class II region genes to enhance antigen presentation and CD4 T-cell activation. This regulation is evidenced by the selection of two upstream variants, **HLA-DRA** (rs9268615) and **HLA-DQA2** (rs9276351), and two downstream variants, **HLA-DQB2** (rs10807113) and **HLA-DMB** (rs3129299). IFN-γ stimulation leads to leukocyte migration (GO:0050900), as evidenced by the selection of two missense variants in **SELL** (rs3177980) and **SELE** (rs5356). SELL (L-selectin) and SELE (E-selectin) initiate leukocyte migration through complementary selectin-mediated mechanisms, forming a multi-step adhesion cascade essential for homing of lymphocytes to secondary lymphoid organs [82]. Furthermore, the humoral immune response process (GO:0006959) is observed, with key regulators including a 3’UTR variant (rs835575) in the **NOTCH2** gene and an intronic variant (rs3780143) in the **PAX5** gene. PAX5 acts as the master transcription factor that commits hematopoietic progenitors to the B lineage in the bone marrow, suppressing non-B genes while activating B-cell-specific programs, such as VDJ recombination and PI3K signalling, to ensure the survival and maturation of follicular, marginal zone, and germinal centre B cells, which are essential for high-affinity antibody production [83]. NOTCH2 signalling diversifies mature B cell fates in the spleen, promoting marginal zone B cells for rapid T-independent responses and influencing germinal centre formation [84]. Complement activation (GO:0006956) is an integral effector and regulatory component of the humoral immune response, involving the upstream variant (rs2921178) in the **C6** component and the synonymous variant (rs12614) in the **CFB** component.

Finally, a recent selection was identified in two variants within metabolic and innate immune-related genes, a downstream variant (rs7622847) in **PPARG** and a missense variant (rs4077515) in **CARD9.** These variants are linked to lipid metabolism, waist-to-hip ratio, and inflammatory bowel diseases, potentially suggesting adaptations to dietary shifts to the local microbiota [85, 86]. However, these signals are often polygenic and subject to complex gene-environment interactions.

**Figure 5:**
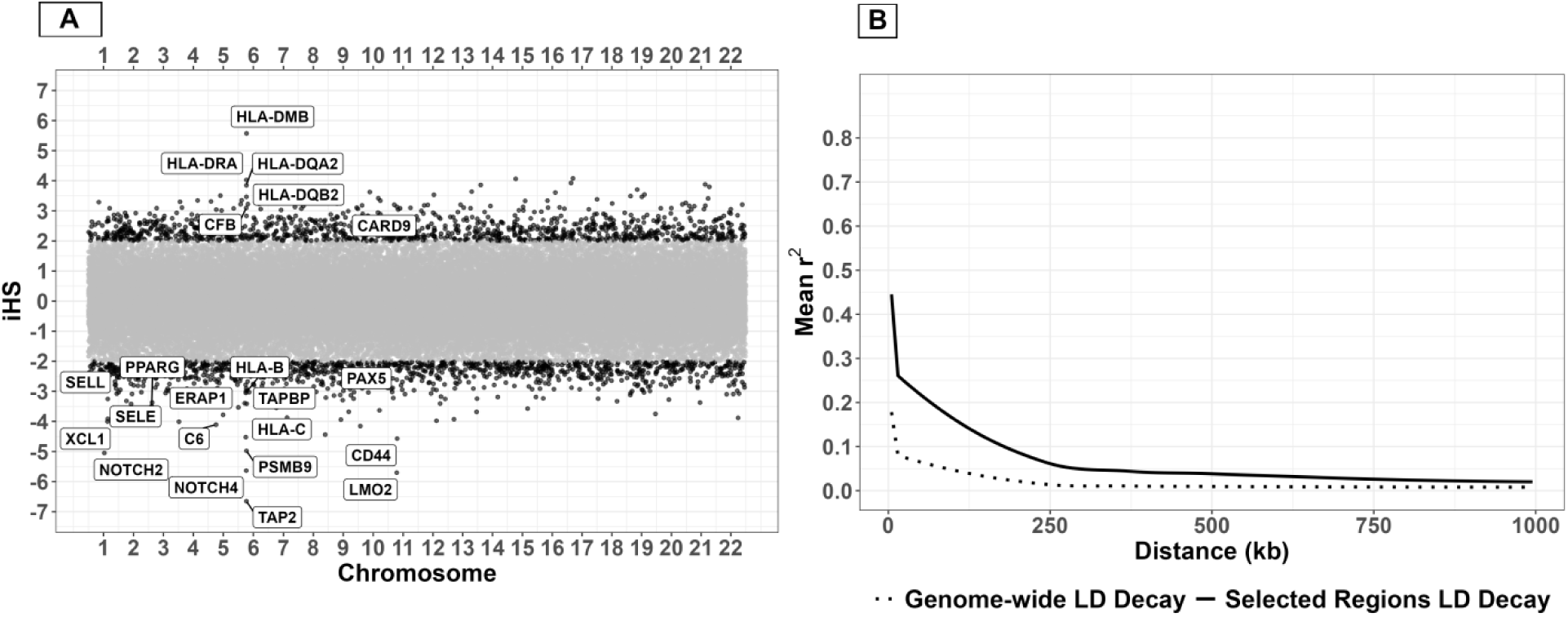
A- Candidate genes under natural selection, identified by genetic variants with elevated iHS values in the Senegalese population. B-LD decay curve for iHS-selected regions is shown in comparison to the genome-wide LD decay curve. The selected regions’ solid curve shows a remarkable difference in mean r² at short distances and slower decay than the dotted genome-wide decay, while remaining substantially elevated above the genome-wide baseline at long distances. The decay pattern is consistent with recent and ongoing selective sweeps.

### PBS-Derived Signature of Pathogen-Driven Immune Adaptation in Thailand

PBS signatures of selection in immune-related genes indicate distinct evolutionary pressures imposed by the region’s historical and contemporary pathogen landscape, including specific adaptations to endemic malaria strains and to bacterial and viral infections. Plasmodium falciparum and Plasmodium vivax exert selective pressure, which have historically been hyperendemic in Southeast Asia, indicating sustained evolutionary pressure from Plasmodium species [87]. We identified two selection signals associated with a downstream gene variant (rs13053958) in **RBFOX2** and a synonymous variant (rs2490741) in **EXOC6**, both of which are involved in erythrocyte differentiation. Furthermore, the selection signal associated with an intronic variant (rs12497256) in the **PXK** gene provides strong evidence of adaptation, given PXK’s direct and indirect roles in red blood cell physiology. PXK encodes a serine/threonine kinase involved in cellular trafficking and signalling, and has been robustly linked to key haematological traits in genome-wide association studies, such as hematocrit and RBC count [88]. These traits are fundamental to oxygen transport and blood viscosity, and their genetic regulation is a well-established target of natural selection in environments characterised by hemolytic stress, particularly those endemic for Plasmodium parasites. Specific PXK variants that influence RBC abundance or haemoglobin content may confer a fitness advantage by optimising oxygen delivery in hot, humid climates or, more importantly, by modifying the RBC intracellular environment to restrict malaria parasite replication and survival [89].

Regarding bacterial infections, the selection signal on a downstream gene variant (rs1407309) in **TNFSF8**, a regulator of CD8+ alpha-beta T-cell differentiation (GO:0043374) and a driver of memory cell formation, suggests T-cell-dependent antimicrobial adaptation against intracellular bacteria prevalent in Thailand, such as Mycobacterium tuberculosis and Treponema [90]. Furthermore, the significant PBS signal for a 3’UTR variant (rs139921) in the **TNRC6B** gene, which participates in the innate immune response and the Fc-epsilon receptor signalling pathway (GO:0038095), indicates IgE-mediated responses essential for defence against soil-transmitted helminths and environmental pathogens, including Burkholderia pseudomallei, the causative agent of melioidosis, which is highly endemic in Northeast Thailand [91]. These naturally selected signals in TNFSF8 and TNRC6B also suggest a dual adaptive role, providing an advantage against endemic viral infections such as papillomaviruses and coronaviruses [92,93]. In addition, the downstream gene variant (rs12980275) in **IFNL3** and the intron variant (rs12979860) in **IFNL4**, both linked to hepatitis viral load and interferon response [94,95], suggest adaptation to endemic hepatotropic viruses prevalent throughout Southeast Asia.

Expanding upon the PBS-identified immune genes, several loci demonstrated broader biogeographical adaptation in the Thai population. The naturally selected missense variant (rs2648307) in **MKRN2** and the intron variant (rs6795441) in **RAF1** are associated with itch intensity following mosquito bites [96]. A reduction in local inflammation and itching sensation may lower the risk of secondary infections from scratching and potentially decrease attractiveness to mosquitoes, thereby conferring a fitness advantage in malaria-endemic regions.

**Figure 6:**
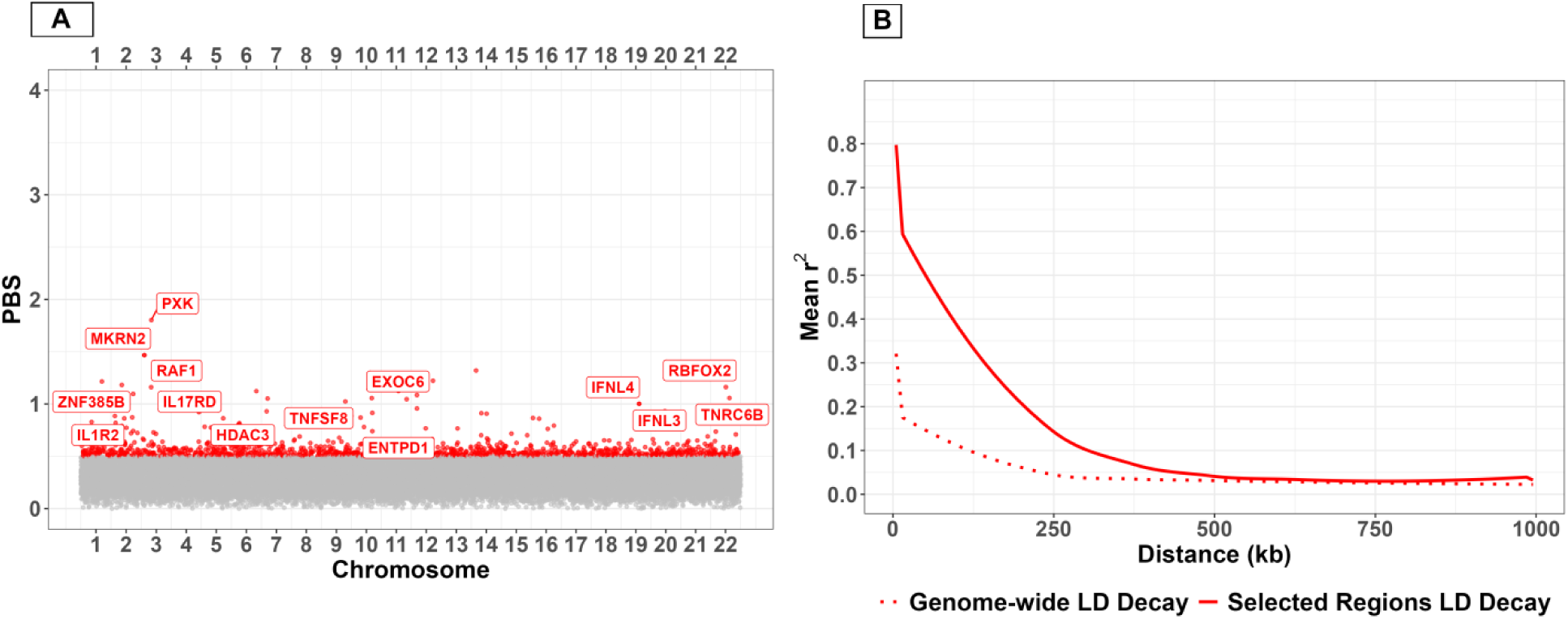
A- Candidate genes under natural selection, identified by genetic variants with elevated PBS values in the Thai population. B-LD decay curve for PBS-selected regions i shown in comparison to the genome-wide LD decay curve. The selected regions’ solid curve shows a remarkable difference in mean r² at short distances and a slower decay than the dotted genome-wide decay. The elevated LD in the selected regions reflects genuine selective sweeps.

### iHS-Derived Signature of Pathogen-Driven Immune Adaptation in Thailand

iHS signatures of recent positive selection in the Thai population closely correspond to the historical and ongoing burden of specific endemic bacterial and viral infections in Southeast Asia [97]. The recent evolutionary history of the Thai population is marked by rapid adaptations at the mucosal barriers driven by enteric viruses and bacteria, alongside continuous fine-tuning of the HLA antigen presentation system. Notably, natural selection targets molecular processes associated with antigen processing and presentation (GO:0019882) via MHC class I (GO:0002474) and MHC class II (GO:0002504), involving a synonymous variant (rs1059510) in **HLA-E**, a missense variant (rs1131896) in **MICA,** an intronic variant (rs2239800) in **HLA-DQA2,** and an upstream gene variant (rs9348894) in **HLA-DQB2**. Furthermore, the selection of the upstream variant (rs12713741) in a kinesin motor protein, **KIF3C**, that contributes to vesicular transport as part of the intracellular machinery for MHC class II molecules from late endosomes to the cell surface, highlights that adaptation can occur not just in the receptors themselves, but in the intracellular machinery required to deploy them [98]. These genes facilitate antigen presentation to CD8+ and CD4+ T cells, and their selection suggests adaptation to intracellular pathogens, including tuberculosis [99], in which HLA class II alleles influence the breadth of the CD4+ T-helper 1 response, and MICA functions as an activating ligand and antigen presentation enhancer. The selection on these genes also reflects adaptation to dengue virus, as specific HLA class I alleles are associated with increased risk of severe dengue, indicating a history of selection for variants that promote a more balanced, less pathogenic T-cell response to this endemic arbovirus [100]. In addition, selection was detected on the intronic variant (rs3917691) in the **SELP** gene. SELP (P-selectin) is critical for modulating inflammation during dengue shock syndrome (DSS) by mediating platelet-leukocyte and platelet-endothelial interactions that exacerbate vascular leakage and cytokine storms [101]. Dengue fever and related arboviruses also exert strong selection on innate immune sensing, particularly the TLR signalling pathways. This includes both MyD88-dependent (GO:0002755) and TRIF-dependent (GO:0035666) signalling, with selection concentrated on the intronic variant (rs1103031) in the kinase gene **RPS6KA2**. Activation of multiple TLRs through MyD88 or TRIF induces type I interferons and pro-inflammatory cytokines, reflecting the necessity for rapid innate detection of viral RNA. Within this viral sensing network, TLR3 signalling (GO:0034138) serves as a primary sensor for viral double-stranded RNA from dengue, Zika, and chikungunya, initiating a critical interferon response to limit early viremia [102]. Similarly, chikungunya virus, another mosquito-borne alphavirus, drives selection on broad innate immune and complement cascades due to its reliance on TLR3-mediated recognition and the essential role of complement in controlling acute arthritogenic viremia [103]. To ensure viral clearance without causing severe immunopathology, a balanced cell-mediated immune response is required. This balance is highlighted by selection on the response to interferon-gamma (GO:0034341), involving a downstream gene variant (rs2745411) in the **UBD** gene. In contrast to the antiviral TLR responses, parallel selection is observed in antibacterial pathways. TLR4 signalling (GO:0034142) is the principal receptor for bacterial lipopolysaccharide, central to recognising and triggering an inflammatory response against typhoid fever and other gram-negative bacterial threats endemic to the region [104].

This recent natural selection signature also corresponds with the hepatitis B (HBV) immune response, where cytotoxic CD8 T cells are essential for eliminating HBV-infected hepatocytes, and specific HLA class I and II alleles determine viral clearance versus progression to chronic infection [105]. In contrast, hepatitis A virus (HAV), which induces lifelong immunity after acute infection, is associated with selection on genes regulating B cell activation (GO:0042113) and humoral immune response (GO:0006959), as evidenced by an intronic variant (rs9819066) in **FOXP1** and another intronic variant (rs16933784) in **PAX5**, supporting the robust humoral response required for HAV resolution. Additionally, T cell-mediated cytotoxicity (GO:0001913) and natural killer cell-mediated cytotoxicity (GO:0042267), involving MICA and HLA-E, are critical for controlling other viral infections, such as Japanese encephalitis virus (JEV) and rabies, in which NK and T cells are essential for clearing virus-infected neurons before fatal neuroinvasion. In neurotropic viral infections like JEV and rabies, T cells concord with neutralising antibodies and innate responses, providing multi-layered immune coordination that may require immune modulation to prevent immunopathology while achieving viral clearance. Notably, a high-impact stop-gained mutation (rs1001420) in the **MROH2B** gene suggests adaptive immune modulation, where gene disruption likely conferred a survival advantage by fine-tuning the immune-inflammatory response. MROH2B loss-of-function aligns with RNAi-mediated knockdown, which decreased NF-κB reporter expression [106], indicating that MROH2B normally acts as a positive regulator of NF-κB signalling. Furthermore, the selection of metabolic genes via a downstream variant (rs4646774) in **ALDH1B1** and a 3’UTR variant (rs11145043) in **FXN** may have been driven by their roles in managing oxidative stress during intense immune responses to infections such as dengue fever or melioidosis, where a robust oxidative burst is necessary but must be tightly regulated to prevent host tissue damage. Concurrently, a strong signal is observed in complement activation (GO:0006956), specifically in the classical (GO:0006958) and alternative (GO:0006957) pathways, involving selection of a missense variant (rs6540433) in **CR2,** an intronic variant (rs2004385) in **C6,** a missense variant (rs13157656) in **C7,** and a synonymous variant (rs700233) in **C9.**

Finally, selection of a missense variant (rs601338) in the **FUT2** gene highlights adaptation at the host-microbe interface. FUT2 (fucosyltransferase 2) encodes an enzyme responsible for secreting ABO histo-blood group antigens onto mucosal surfaces of the gastrointestinal and respiratory tracts. Functional variation in FUT2, which influences the glycan landscape, is a key determinant of host susceptibility to a range of pathogens. Certain FUT2 polymorphisms that modify antigen presentation are associated with differential resistance to norovirus and Helicobacter pylori, and may affect the risk of enteric infections such as cholera, which has a historical presence in Southeast Asia [107,108]. In Thailand, where diarrheal and foodborne illnesses remain persistent public health challenges, selection on functional FUT2 alleles likely reflects evolutionary optimisation of the mucosal glycome to resist colonisation or invasion by prevalent enteric pathogens, while also shaping gut microbiota composition to enhance metabolic efficiency and immune priming [109]. Supporting the theme of barrier defence, we identified a selected upstream variant (rs7302661) in **IL22**, a cytokine essential for maintaining the integrity of the intestinal and respiratory barriers and secreting antimicrobial peptides [110]. In addition, a missense variant(rs5741804) in the **BPI** gene, also known as bactericidal/permeability-increasing protein, is expressed by epithelial cells and specifically targets and neutralises the lipopolysaccharide of gram-negative bacteria [111].

**Figure 7:**
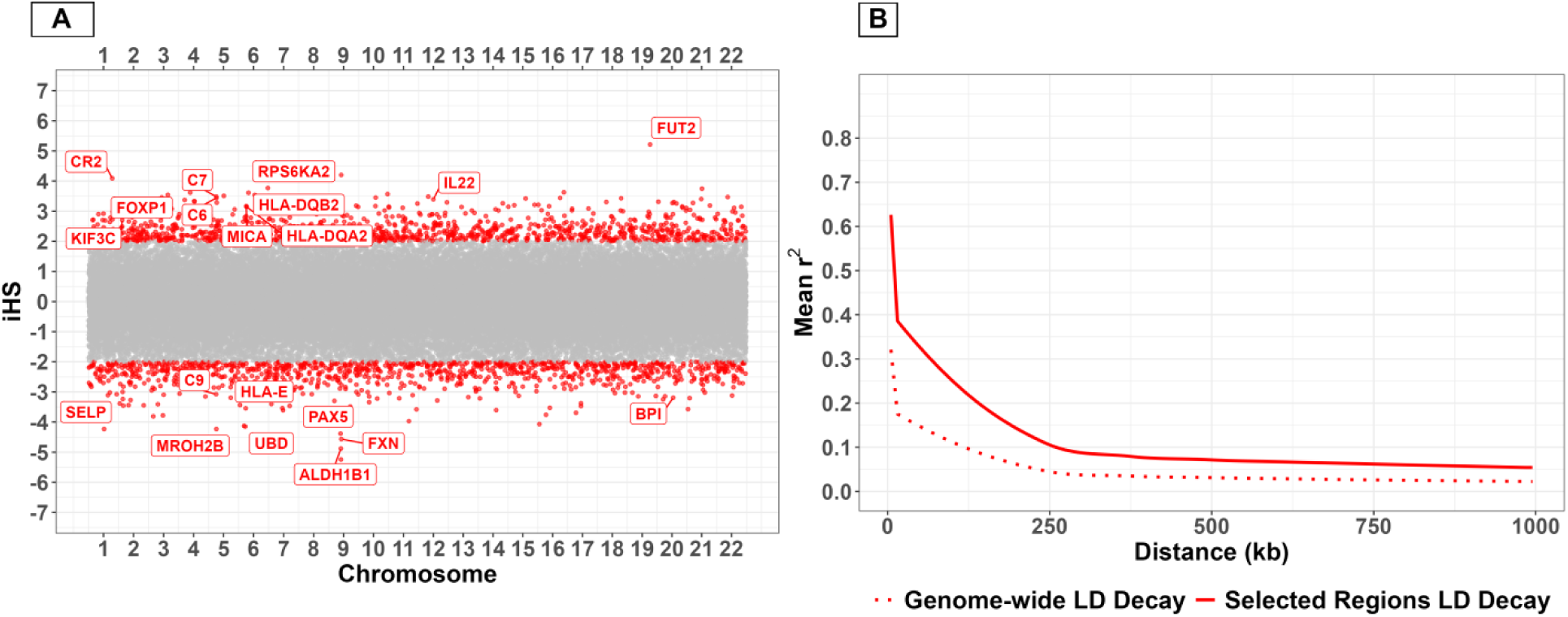
A-Candidate genes under natural selection, identified by genetic variants with elevated iHS values in the Thai population. B-LD decay curve for iHS-selected regions i shown in comparison to the genome-wide LD decay curve. The selected regions’ solid curve shows a remarkable difference in mean r² at short distances and slower decay than the dotted genome-wide decay, while remaining substantially elevated above the genome-wide baseline at long distances. The decay pattern is consistent with recent and ongoing selective sweeps.

### PBS-Derived Signature of Pathogen-Driven Immune Adaptation in Peru

PBS signatures of selection identified in Peruvian populations provide robust genomic evidence of sustained adaptation to a high-pathogen environment, with a particularly strong signature of defence against endemic helminths. This finding is supported by functional annotation of the top candidate loci, which reveals genes and pathways specifically associated with Type 2 immunity. This type of immunity is the primary host defence mechanism against metazoan parasites, characterised by IgE production and eosinophil activation [112]. The most direct evidence is the annotation of the Fc-epsilon receptor signalling pathway (GO:0038095), which involves selecting an intronic variant (rs12884681) in **IGHE**. The IGHE gene encodes Immunoglobulin E (IgE), the key antibody isotype in anti-helminthic immunity, which cross-links the Fc-epsilon receptor on eosinophils and mast cells, triggering degranulation and parasite damage [113]. Th evolutionary refinement of this pathway is further supported by the co-selection of related processes, such as eosinophil chemotaxis mediated by the intronic variant (rs2171544) in the **HRH1** gene. HRH1, the histamine H1 receptor on eosinophils, enhances their activation by promoting shape changes, adhesion, and chemotaxis priming, fine-tuning histamine-mediated boosts during Type 2 inflammation. Complementing this, the synonymous variant (rs2073342) in the **RNASE3** gene, also known as eosinophil cationic protein, which acts as the cytotoxic effector, is released from eosinophil granules to disrupt helminth cuticles [114]. This anti-helminthic genetic signature is substantiated by paleoparasitological studies in the Americas, which confirm a high prevalence of soil-transmitted helminths such as Trichuris trichiura and Ascaris lumbricoides, thereby establishing persistent selective pressure [115]. This adaptation likely entailed an evolutionary trade-off as the genetic variants conferring anti-helminthic immunity may have concurrently increased susceptibility to atopic disorders, including asthma, in contemporary populations. This phenomenon is described as the “hygiene hypothesis” in reverse, in which strong selection for Th2 responses predisposes individuals to hypersensitivity [116]. The Fc-epsilon receptor signalling pathway, when not engaged by helminths, can become hyperresponsive to harmless environmental allergens, leading to mast cell and basophil degranulation. Additionally, eosinophil chemotaxis is a hallmark of the inflammatory pathology observed in asthma and other atopic diseases, in which eosinophils cause tissue damage in the airways and skin [117]. This genetic background may contribute to Peru’s high asthma burden, with prevalence estimates reaching 19.6% in Lima [118]. Consequently, selection has also acted on genes regulating immune balance, including an intronic variant (rs202654) in the **TOB2** gene. TOB2 is a transcriptional regulator that suppresses T cells and TLR pathways [119]. These variants appear to have been selected to modulate inflammatory responses, preventing excessive tissue damage during chronic helminthiasis, while also contributing to the increased risk of inflammatory and metabolic disorders in modern populations [120].

The Peruvian adaptation is not limited to a single pathogen. Instead, it reflects a broad evolutionary refinement of immunity to address diverse and persistent endemic threats, including mycobacterial infections such as tuberculosis [121] and viral infections such as HBV [122]. This adaptation is characterised by selective pressure on key regulatory hub genes that coordinate both innate recognition and adaptive effector mechanisms. In particular, the splice polypyrimidine tract variant (rs2306696) affects the hub gene **IRAK2**, which is central to multiple TLR signalling pathways, including TLR2 signalling (GO:0034134), and represents a critical evolutionary target for enhancing innate immune sensing [123]. Additionally, a missense variant (rs20551) in the hub gene **EP300**, a transcriptional co-activator, integrates NF-κB, STAT6/GATA3, and Foxp3 signalling during bacterial infections [124–126]. This selection signature suggests anti-bacterial preparedness through neutrophil orchestration and epithelial protection. As per the selection of an upstream gene variant (rs17322627) in **CMKLR1**, a chemerin receptor on epithelial cells, promotes epithelial antimicrobial defence by upregulating lactoperoxidase, which generates antimicrobial hypothiocyanite and helps restrict Gram-negative bacterial overgrowth in the gut and skin mucosa [127]. In addition, dominant selection occurs within the CXC chemokine cluster, specifically two 3’UTR variants in **CXCL5** (rs3775488) and **CXCL2** (rs9131). CXCL5 and CXCL2, secreted by epithelial cells and macrophages during pulmonary or tissue infection, act as potent neutrophil chemoattractants via CXCR2, establishing chemotactic gradients that drive rapid influx and engulfment of extracellular bacteria [128,129]. The innate signalling cascade primes the adaptive response, as evidenced by selection of a synonymous variant (rs4606515) in the hub gene **CD3E.** The **CD3E** gene is essential for T cell receptor signalling (GO:0050852) and related processes such as T cell costimulation (GO:0031295), activation (GO:0042110), and positive regulation of T cell proliferation (GO:0042102). Efficient T-cell immunity, particularly CD8+ cytotoxic T-cell function, is crucial for controlling intracellular pathogens [130]. Selection on CD3E indicates evolutionary fine-tuning of cellular immunity to enhance pathogen clearance, a finding supported by population genomic studies linking T-cell signalling genes to pathogen-driven selection in humans. Furthermore, a strong selection signal is observed for an intronic variant (rs4308217) in the **CD86** gene, a key co-stimulatory molecule on antigen-presenting cells that ensures robust T-cell activation and adaptive immunity. Evolutionary analyses show that CD86 and other costimulatory genes have been targets of positive selection, consistent with pathogen-driven pressures that balance effective T-cell immunity against increased risk of autoimmunity [131]. Together with an intronic variant (rs2735835) in the **CCL18** gene, which is expressed by macrophages and recruits regulatory T cells [132]. In contrast, a downstream gene variant (rs7283760) in the **ICOSLG** gene, which encodes the inducible T-cell costimulator ligand, is pivotal for T-cell activation, proliferation, and germinal centre formation, which are necessary for B-cell antibody production [133], and its selection is likely to stabilise adaptive immune responses against tuberculosis. In addition to the selection of an intronic variant in the **CXCR5** gene. CXCR5 is the receptor for CXCL13 and is central for organising B cell areas, germinal centres, and T-cell help that generate effective antibody responses [134]. Finally, selection of an intronic variant (rs12656176) in the hub gene **PIK3R1** highlights its role as a central integrator, involved in Fc-gamma receptor signalling during phagocytosis (GO:0038096), T cell receptor signalling (GO:0050852), and B-cell differentiation (GO:0030183), supporting humoral responses to viruses. This coordinated selection on innate IRAK2, signalling PIK3R1, transcriptional EP300, and adaptive CD3E immune hubs suggests evolutionary adaptation across pathogen sensing, cellular activation, and effector function. This genetic architecture provided the Andean populations with a robust, balanced defence capable of confronting the polymicrobial landscape of ancient Peru, where respiratory pathogens such as tuberculosis and endemic viruses posed continuous threats to survival and reproductive fitness.

**Figure 8:**
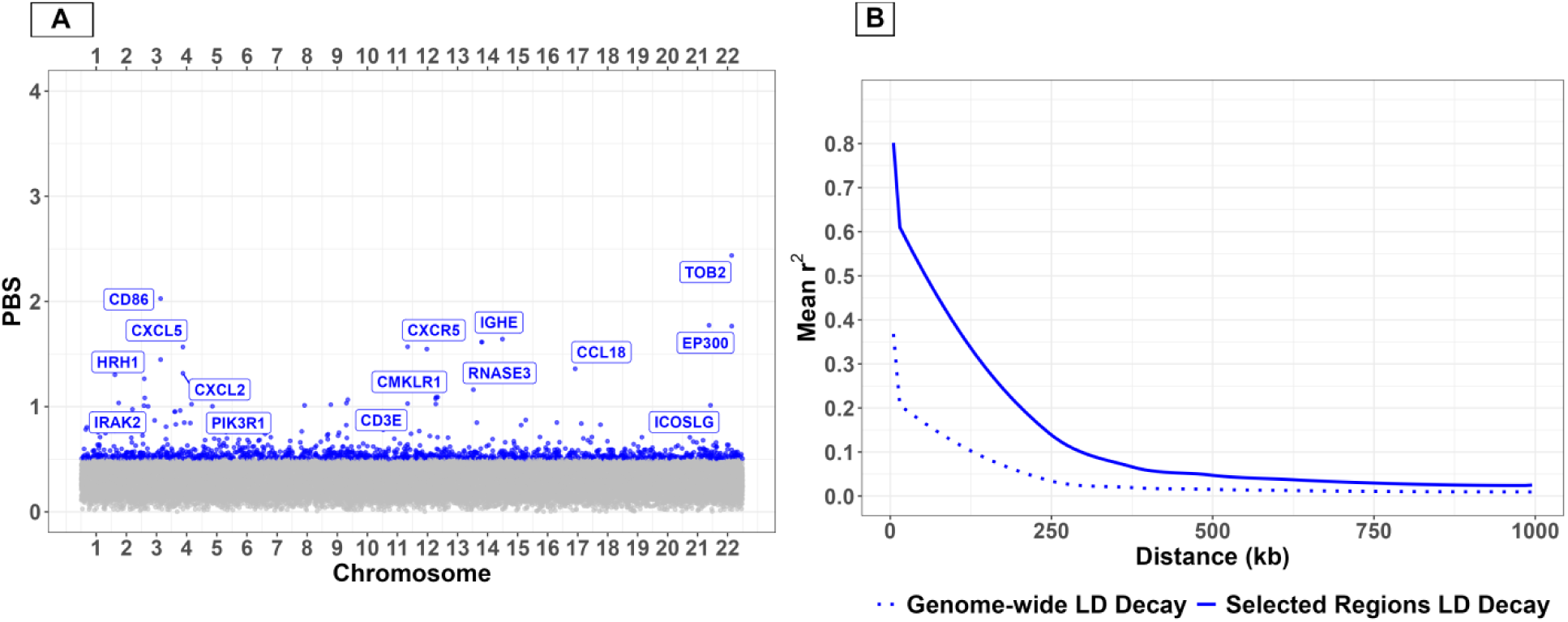
A-Candidate genes under natural selection, identified by genetic variants with elevated PBS values in the Peruvian population. B-LD decay curve for PBS-selected regions is shown in comparison to the genome-wide LD decay curve. The selected regions’ solid curve shows a remarkable difference in mean r² at short distances and a slower decay than the dotted genome-wide decay. The elevated LD in the selected regions reflects genuine selective sweeps.

### iHS-Derived Signature of Pathogen-Driven Immune Adaptation in Peru

iHS signatures of recent positive selection in the Peruvian population align with the ongoing burden of specific endemic infectious diseases in the region and includes evidence of recent adaptation to helminths, as evidenced by the selection of the missense variant (rs2256721) in the **CHIA** gene, which contributes to the immune response to helminths by modulating IgE-mediated type-2 immunity, as genetic variants in CHIA are associated with the intensity of IgE responses to helminth antigens [135]. In addition, the selection of the synonymous variant (rs2306888) in the **TGFBR3** gene, which encodes a co-receptor for the TGF-β signalling pathway that helminths exploit to induce regulatory T cells and modulate effector immunity [136].

The immune defence architecture is centred around the interferon signalling axis, encompassing response to type I interferon (GO:0034340) and response to interferon gamma (GO:0034341), together with their positive regulation represented by the two processes: positive regulation of type I interferon-mediated signalling pathway (GO:0060340) and positive regulation of interferon gamma-mediated signalling pathway (GO:0060335). Functional annotation revealed these processes following selection of an upstream variant (rs3754944) in the **SP100** gene and a synonymous variant (rs13339199) in the **NLRC5** gene. Type I IFN-α/β signalling induces a generalised antiviral state, crucial for limiting replication upon exposure to endemic arboviruses such as Dengue and Zika [137], while type II IFN-γ activates macrophages to control intracellular pathogens such as Mycobacterium tuberculosis [138] and Bartonella bacilliformis [139]. Both types IFN-α/β and IFN-γ responses strongly upregulate MHC class I antigen processing and presentation pathways to enhance CD8+ T cell recognition of infected or transformed cells, including cellular response to interferon gamma (GO:0071346), antigen processing and presentation of peptide antigen via MHC class I (GO:0002474), peptide antigen transport (GO:0046968), and positive regulation of T cell-mediated cytotoxicity (GO:0001916). These processes are primarily associated with the selection of a missense variant (rs2230301) in the **EPRS** gene and an intronic variant (rs241425) in the **TAP2** gene, and these are considered fundamental processes for immune surveillance against infected cells, serving as a critical defence mechanism against the endemic viral and bacterial infections and also against the intracellular phases of parasites such as Chagas disease [140]. At barrier sites, innate mucosal and effector mechanisms, such as the innate immune response in mucosa (GO:0002227) and antibacterial peptide secretion (GO:0002779), involving a missense variant (rs10502001) in the **MMP7** gene and two intronic variants in the nitric oxide synthase genes **NOS1** (rs7133438) and **NOS2** (rs3730013). The NOS2 gene provides immediate protection against intracellular bacteria such as Mycobacterium tuberculosis [141]. However, nitric oxide is also a critical vasodilator [142]. In the context of high-altitude hypoxia in the Andes, altered nitric oxide production is a known physiological adaptation to regulate pulmonary blood pressure and oxygen delivery. Therefore, the simultaneous selection on multiple NOS genes suggests a dual-purpose adaptation: balancing pathogen defence with high-altitude cardiovascular homeostasis.

Further structuring the immune response are leukocyte migration and chemotaxis, including positive regulation of macrophage chemotaxis (GO:0010759) via an intronic variant (rs4964676) in **CMKLR1** and neutrophil chemotaxis (GO:0030593) via an intronic variant (rs11750248) in **ITGA1**. Selection of an intronic variant (rs548175) in the **MAML2** gene, a transcriptional co-activator in the Notch signalling pathway, is essential for lymphocyte development and may influence the formation of the adaptive immune repertoire [143]. For sustained immunity and pathogen clearance, the humoral and B-cell-mediated arm is highlighted by B cell activation (GO:0042113), B cell receptor signalling pathway (GO:0050853), and humoral immune response (GO:0006959), driven by a 5’UTR variant (rs12506517) in **BANK1,** an intronic variant (rs8096471) in **BCL2**, and an intronic variant (rs17391260) in **PAX5**. This arm of the immune response is essential for generating neutralising antibodies against circulating arboviruses and facilitating phagocytic clearance of bacterial infections. Finally, regulatory and homeostatic processes such as T cell homeostasis (GO:0043029) and B cell homeostasis (GO:0001782) ensure a balanced adaptive response, preventing immunopathology while maintaining the capacity for cytotoxic clearance of infected cells. This function is vital for controlling chronic infections such as Tuberculosis and Chagas disease, both prevalent in the region.

In summary, this genetic architecture—characterised by enhanced interferon responses, potent antigen presentation, regulated leukocyte trafficking, and balanced B and T cell immunity— demonstrates population-specific immunological fine-tuning shaped by the cumulative burden of Peru’s distinctive endemic pathogen landscape.

**Figure 9:**
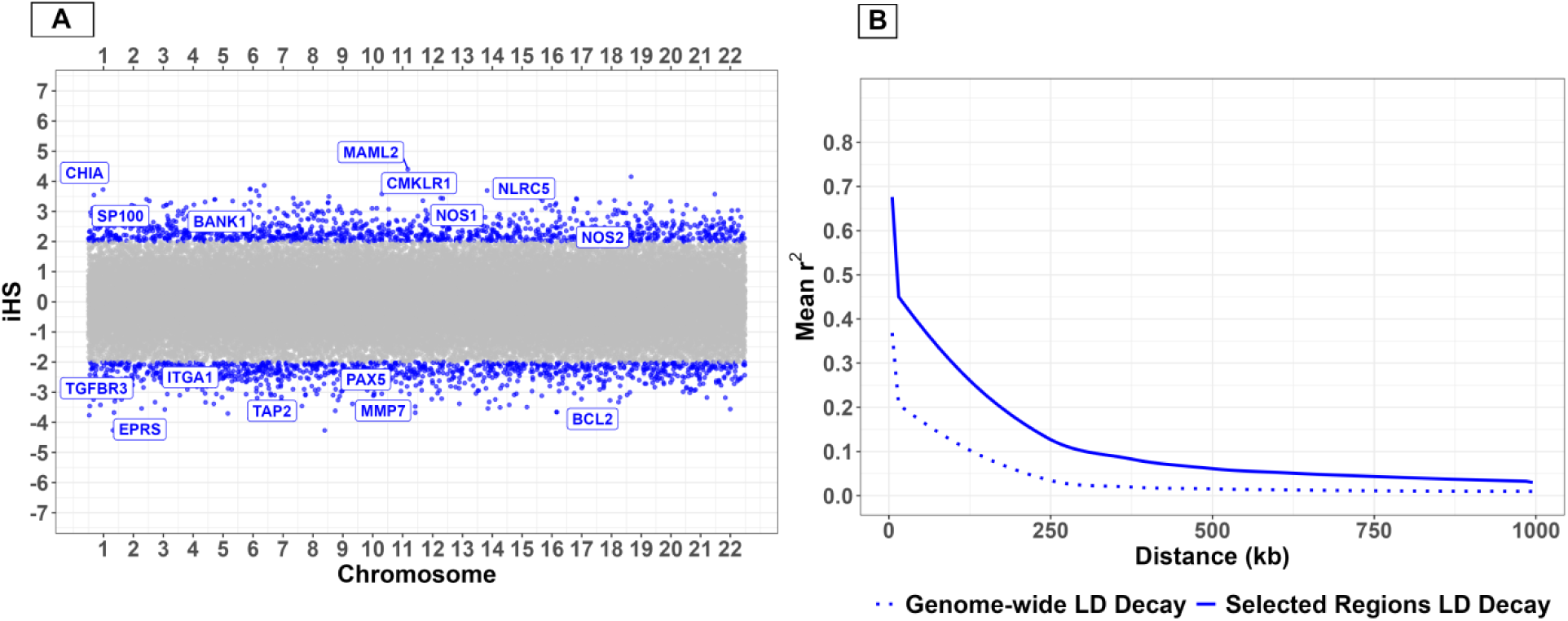
A-Candidate genes under natural selection, identified by genetic variants with elevated iHS values in the Peruvian population. B-LD decay curve for iHS-selected regions is shown in comparison to the genome-wide LD decay curve. The selected regions’ solid curve shows a remarkable difference in mean r² at short distances and slower decay than the dotted genome-wide decay, while remaining substantially elevated above the genome-wide baseline at long distances. The decay pattern is consistent with recent and ongoing selective sweeps.

## Discussion

Over millennia, distinct infectious pressures have left genetic footprints in human populations. By integrating PBS, which reflects deep population-differentiating selection, and iHS, which captures recent, population-specific sweeps, this study demonstrates that the immune-related genetic architecture of Senegal, Thailand, and Peru has likely been durably shaped by each region’s unique pathogenic landscapes. Genetic variants under selection in immune-related genes are directly relevant to immune response research and may inform theoretical frameworks for ancestry-informed vaccine design by highlighting variants that influence antigen presentation, interferon signalling, hematopoiesis, and mucosal immunity. Alleles under selection in antigen presentation and HLA genes, which vary significantly across ancestral populations, may influence which pathogen epitopes are presented to adaptive immune cells, thereby potentially modulating immunogenicity and guiding future epitope selection strategies [144]. Genes in the innate immune and interferon pathways are central to infection sensing and the amplification of vaccine-induced responses. Understanding ancestral variation in these genes could enable the exploration of adjuvants that optimally engage these selected pathways [145]. Furthermore, immune regulatory pathways may influence the magnitude and durability of vaccine-induced responses by modulating early innate activation and the establishment of long-lived adaptive immunity [146]. Hematopoiesis genes are also relevant for vaccines that require rapid expansion of antigen-specific B and T cells, as inherited differences in hematopoietic regulation may affect baseline immune cell composition and proliferative capacity, thereby influencing the magnitude and speed of vaccine-elicited immune responses [147]. Finally, signatures in mucosal immunity genes might predict the theoretical efficacy of orally administered vaccines by reflecting population-specific host adaptations to recurrent local mucosal pathogens [148].

In Senegal, Plasmodium falciparum malaria appears to have exerted the predominant selective pressure. Genetic variants under selection include those involved in erythrocyte development and differentiation, specifically ANK1, ID2, and FOXO3, as well as RBFOX2 and EXOC6, which govern cytoskeletal maturation and lineage commitment, both of which are essential for malaria resilience. Selection of ITPR3 highlights Fcγ receptor-mediated phagocytosis, while TLR4 and TRAF6 are involved in toll-like receptor signalling. Complement activation is also integral, evidenced by selection on MASP1, C4BPA, and C4BPB. These findings provide hypothesis-generating insights into the development of malaria and tuberculosis vaccines. For malaria vaccines, the selection of C4BPA is particularly relevant because the oligomerisation domain of the α-chain of C4b-binding protein has been shown to enhance T cell responses when fused to Plasmodium antigens in adenoviral vectored vaccines [149]. Strong selection in the complement classical and lectin pathways, alongside Fcγ receptor signalling, suggests a hypothesis that malaria vaccines might benefit from eliciting antibodies capable of efficient complement fixation and Fcγ receptor-mediated opsonic phagocytosis, potentially enhancing parasite clearance [150]. For tuberculosis vaccines, strong selection for TLR4 and TRAF6, which are central to mycobacterial recognition and innate signalling, suggests that adjuvants engaging these pathways could theoretically enhance protective immunity. Tuberculosis vaccine candidates utilising TLR4 agonists within the adjuvant system might activate TRAF6-dependent signalling pathways, potentially limiting early tuberculosis replication and reducing the risk of progression to active pulmonary disease. Selection on NOTCH4, a negative regulator of mycobacteria-induced inflammation, indicates population-specific variation in susceptibility to and severity of tuberculosis, consistent with a theoretical role in shaping tuberculosis immunopathology [151]. Senegalese immune adaptation also offers theoretical insights for HBV vaccine development. Selection on MHC class I antigen-processing genes, including HLA-B, HLA-C, TAP2, and ERAP1, as well as on immunoproteasomes such as PSMB9 and CTL-priming chemokines like XCL1, reflects adaptation to endemic HBV infections. This selection represents candidate pathways for therapeutic cytotoxic vaccines, as viral-vectored vaccines delivering antigens into the host cell cytoplasm might exploit the enhanced ERAP1/TAP2 peptide-loading complex [152]. Prophylactic HBV vaccines could also be explored in this context due to selection on MHC class II antigen-presenting genes, specifically HLA-DQA2 and HLA-DQB2, B-cell commitment genes such as PAX5 and NOTCH2, and regulatory genes including BCL6, BCL2L15, DHX58, and PTPN22, further suggesting ancestry-specific humoral immunity architectures [153]. Vaccine pharmacogenomics may therefore provide theoretical frameworks for designing HBV vaccines optimised for both humoral and cellular immunity in this population.

In Thailand, signatures of selection indicate adaptation to a polymicrobial environment, with immune architecture likely influenced by malaria, arboviral, hepatotropic viral, and bacterial pressures. Selection signals include variants in the interferon-λ genes IFNL3 and IFNL4, as well as the antigen presentation genes HLA-E, MICA, HLA-DQA2, and HLA-DQB2. Selection of IFNL3 and IFNL4 variants may influence hepatitis virus clearance and vaccine response, as patients with altered immunocompetence exhibit variation in their immunological response to hepatotropic viruses [154]. Selection signatures in B-cell commitment genes, specifically FOXP1 and PAX5, support an evolutionary contribution of B-cell developmental pathways to humoral vaccine responsiveness [155], while selection of MHC class I alleles suggests that CD8+ T cell epitope selection for arboviral vaccines might need to account for common HLA variants in Thai populations. Selection of innate viral RNA sensors, such as RPS6KA2 and UBD, supports the hypothesis that arboviral vaccines could achieve higher efficacy by utilising TLR3-agonist adjuvants to activate the population’s adapted type I interferon pathways [156,157]. The selection of the SELP gene, which modulates vascular leakage during dengue shock syndrome, suggests a need for careful evaluation of live attenuated vaccines in this population to assess potential effects on vascular leakage and endothelial activation. Selection of FUT2 is critical for oral vaccines, as FUT2 determines the expression of ABO histo-blood group antigens on mucosal surfaces. These antigens serve as receptors for enteric pathogens and affect susceptibility to norovirus and rotavirus, which may directly impact the administration and immunogenicity of live-attenuated oral vaccines. Thai adaptation against bacterial threats is associated with selection signals in the TNFSF8 gene, which contributes to the generation of CD8+ memory T cells, as well as in the MHC class II alleles HLA-DQA2 and HLA-DQB2. These findings suggest a theoretical framework for exploring tuberculosis vaccines to maximise a broad and durable T-cell memory network [158].

In Peru, soil-transmitted helminths appear to have exerted the predominant selective pressure. The Peruvian genome exhibits strong selection on Type 2 immunity genes, including the Fc-epsilon receptor signalling component IGHE, the eosinophil chemotaxis regulators HRH1 and CXCL5, and the immune modulators CCL18 and TOB2. Selection on the innate immune hubs IRAK2, EP300, and PIK3R1, alongside the T-cell activation genes CD3E and CD86, likely reflects adaptation to endemic tuberculosis and arboviruses. Recent iHS signals indicate selection on the interferon response genes NLRC5 and SP100, as well as the antigen processing component TAP2, consistent with ongoing viral pressure. Additional selection in genes involved in B-cell activation and homeostasis, namely BANK1, BCL2, and PAX5, suggests an evolving humoral response against endemic arboviral threats. Selection of TGFBR3 points to ongoing pressure from parasites, highlighting the potential theoretical value of tailored blood-stage vaccine approaches [159]. In helminth-endemic regions, the chronic Th2-biased immune profile may theoretically compromise vaccine responses that require Th1-type cellular immunity. Experimental evidence demonstrates that helminth infections induce a dominant Th2 cytokine profile and impair Th1 responses [160]. Eradication of helminth infections has been shown to reverse these effects, suggesting a hypothesis that deworming prior to immunisation and the use of Th2-modulating adjuvants might be explored to achieve protective cellular immune responses. Selection on MMP7 and NOS2 highlights a robust innate mucosal defence system, suggesting that vaccine strategies engaging these pathways via mucosal routes and Th2-modulating adjuvants might better exploit this evolved mucosal barrier to achieve improved protection [161].

In summary, the development of ancestry-informed vaccines represents a theoretical framework for health equity, aiming to incorporate genetic diversity into standard vaccine formulations to explore equitable protection across global populations. Population-tuned pathways provide hypothesis-generating support for the exploration of ancestry-informed vaccines with population-specific optimisations, potentially facilitating the development of broadly effective and durable vaccines across ancestries. All three populations exhibit selection in erythropoiesis genes, supporting the theoretical exploration of malaria blood-stage vaccine candidates with population-specific adjuvants. Peru may require the exploration of strong Th1-polarising adjuvants to counteract its helminth-driven Th2 baseline. In contrast, Thailand’s selection in TLR3 signalling and Senegal’s in MHC class I effector machinery indicate fundamentally different theoretical arboviral vaccine designs, mainly exploring TLR3-agonist adjuvanted virus-like particles for Thailand and viral-vectored platforms for Senegal. Divergent adaptations in mucosal immunity, such as FUT2 in Thailand and MMP7 and NOS2 in Peru, suggest that oral and mucosal vaccine delivery platforms could be tailored to regional architectures.

A further strength of this comparative analysis is that it reveals shared themes of natural selection across all three populations. First, antigen presentation emerges as a recurring target, especially through HLA-related and peptide-processing genes, reflecting the importance of cytotoxic and helper T-cell surveillance in diverse infectious settings. Second, the interferon signalling axis is universally targeted, though via different nodes: regulatory genes like DHX58 in Senegal, specific cytokines like IFNL3 and IFNL4 in Thailand, and downstream effectors like SP100 and NLRC5 in Peru. Third, immune regulation is repeatedly selected alongside immune activation, indicating that activated immunity is constrained by the need to prevent immunopathology. Fourth, both innate and adaptive arms are involved in all three populations, and adaptive immunity involves cytotoxic T cells and B cell antibodies, refuting a simplistic model in which adaptation is driven by only one branch of immunity. Instead, the data support a layered architecture in which selection is driven by sensing, multiple effector, and regulatory processes simultaneously.

## Conclusion

This comparative population genomics study demonstrates that the combined application of PBS and iHS can reveal both shared and population-specific signatures of immune adaptation shaped by distinct regional pathogen landscapes. By identifying immune pathways recurrently targeted by natural selection, our findings offer a theoretical foundation for exploring multiple ancestry-informed vaccine strategies, including antigen selection, adjuvant design, and delivery platforms tailored to population-specific immune architectures. We propose that this evolutionary genomics approach can guide future experimental vaccinology studies and contribute to more equitable, globally attuned vaccine development.

## Supplementary Data

Tables of selected LD blocks with PBS > 0.5 for the three populations, highlighting target genes and their associated GO terms.

Tables of selected LD blocks with |iHS| > 2 for the three populations, highlighting target genes and their associated GO terms.

## Availability of data and materials

The datasets used and analysed during the current study are available from the corresponding author on reasonable request.

## Author Contributions

A.S. was involved in planning, supervising the work, and providing feedback. K.V.S. verified the analytical methods, provided feedback, and helped shape the analysis and manuscript. A.T. planned and performed the analysis, drafted the manuscript, and designed the figures. All authors have read and agreed to the published version of the manuscript.

